# The Pebble/Rho1/Anillin pathway controls polyploidization and axonal wrapping activity in the glial cells of the *Drosophila* eye

**DOI:** 10.1101/2020.02.12.945600

**Authors:** Lígia Tavares, Patrícia Grácio, Raquel Ramos, Rui Traquete, João B. Relvas, Paulo S. Pereira

## Abstract

During development glial cell are crucially important for the establishment of neuronal networks. Proliferation and migration of glial cells can be modulated by neurons, and in turn glial cells can differentiate to assume key roles such as axonal wrapping and targeting. To explore the roles of actin cytoskeletal rearrangements in glial cells, we studied the function of Rho1 in *Drosophila* developing visual system. We show that the Pebble (RhoGEF)/Rho1/Anillin pathway is required for glia proliferation and to prevent the formation of large polyploid perineurial glial cells, which can still migrate into the eye disc if generated. Surprisingly, this Rho1 pathway is not necessary to establish the total glial membrane area or for the differentiation of the polyploid perineurial cells. The resulting polyploid wrapping glial cells are able to initiate wrapping of axons in the basal eye disc, however the arrangement and density of glia nuclei and membrane processes in the optic stalk are altered and the ensheathing of the photoreceptor axonal fascicles is reduced.

## Introduction

In the context of a growing nervous system, neurons, glia, and their precursor cells tightly coordinate cell proliferation and migration. During *Drosophila* larval eye development, perineurial glia (the outermost glia subtype) migrate from the optic lobe towards the optic stalk and the basal surface of the eye disc, along the two subperineurial glial cells (Silies et al., 2007). Migration of perineurial glia into the eye disc is abrogated if the expression of the αPS integrin heterodimer is repressed, causing a loss of focal adhesions and a reduction in the stiffness of the extracellular matrix layer (neural lamella) (Kim et al., 2014; Tavares et al., 2015; Xie et al., 2014). Once perineurial glia contacts the differentiating photoreceptor axons they differentiate into wrapping glia and extend their processes back toward the CNS, ensheathing the axonal fascicles (Hummel et al., 2002; Silies et al., 2007; Tavares et al., 2015). During the growth of the nervous system perineurial glia proliferate extensively (Avet-Rochex et al., 2012; Awasaki et al., 2008), and their cell number in the eye disc is positively controlled by the FGF/Pyramus and Dpp pathways, and by the cell autonomous functions of the Yorkie/Yap, Myc, Dref, and Pdm3 transcription factors (Bauke et al., 2015; Rangarajan et al., 2001; Reddy and Irvine, 2011; Tavares et al., 2017; Yuva-Aydemir et al., 2011). In contrast, subperineurial glial cells increase their individual cell size by endoreplication, undergoing both endocycles and endomitosis, and the increased surface area allows for a correct formation of the blood-brain barrier (BBB) (Unhavaithaya and Orr-Weaver, 2012; Von Stetina et al., 2018; Zulbahar et al., 2018). Glia migration and differentiation are tightly regulated by photoreceptor precursors fitness and differentiation states (Silies et al., 2010; Tavares et al., 2017). In particular, the transition between the perineurial and wrapping glial cell fate is under the control of Thisbe, a FGF8-like ligand that is expressed by differentiating photoreceptors (Franzdottir et al., 2009). This controlled process allows wrapping glia to perform the ensheathment of newly differentiated photoreceptor axons that target the optic lobe (Chang et al., 2018; Chen et al., 2017; Silies et al., 2010). There, wrapping glia also has a role in promoting the differentiation of lamina neurons by secreting Insulin-like peptides in response to EGF secreted from photoreceptors (Fernandes VM, 2017).

By regulating the actin cytoskeleton, the Rho GTPases (RhoA, B and C in vertebrates, and Rho1 in *Drosophila*) have key roles in multiple processes, including cell migration and epithelial morphogenesis, mitosis and cytokinesis (Etienne-Manneville and Hall, 2002; Haga and Ridley, 2016; Raftopoulou and Hall, 2004; Van Aelst and Symons, 2002). However their functions in glial cells are yet poorly characterized. RhoA was shown to promote *in vitro* proliferation of PNS Schwann cells through AKT activation (Tan et al., 2018), and differentiation through the repression of JNK pathway, and both roles were independent of its downstream effector ROCK (Wen et al., 2018). In contrast, RhoA activity inhibited differentiation of oligodendrocytes (Liang et al., 2004) and rescued proliferation of oligodendrocyte precursor cells in *gpr56^stl13/stl13^* mutant zebrafish (Ackerman et al., 2015). In the *Drosophila* PNS, Rho1 was shown to control actin rearrangement as overexpression of constitutively-active Rho1^V14^ in glia caused stalling of cell bodies and inhibited peripheral extension of cell processes (Sepp and Auld, 2003a, b). However, glial expression of dominant-negative Rho1^N19^ (Rho1^DN^) in peripheral glia caused minor glial and axonal phenotypes (Sepp and Auld, 2003a, b), preventing a more complete characterization of the functions of Rho1 in glia-neuron crosstalk. In here, we used the *Drosophila* eye imaginal disc to address the role of Rho1 in retinal glia development and ommatidial axon wrapping. We show that glia-targeted Rho1 knockdown led to perineurial glia migration and proliferation defects. Surprisingly, perineurial glia became more adaptable, displaying an increased ploidy and cell membrane area, maintaining the ability to cover the photoreceptor differentiated domain. However, even though perineurial glial cells initiate differentiation and axonal wrapping in the eye disc proper, in the optic stalk Rho1 repression blocks complete wrapping of 8-axons ommatidial fascicle.

### Repression of the Pebble/Rho1/Anillin genetic pathway in the eye disc glia causes a switch from mitosis to endoreplication

To determine if Rho1 regulates glia development in the developing *Drosophila* eye (Fig. 1A– C), we expressed both Rho RNAi and dominant-negative Rho1 (Rho1^DN^) using the pan-glial *repo*-Gal4 driver (Fig. 1D-F’). Rho1 loss-of-function (LOF) induced a dramatic reduction in retinal glia numbers in third-instar eye discs (from 147 to 29, n=10-19 eye discs, p<0.0001) (Fig. 1G). The reduction of Rho1 function did not interfere with the dispersion of glial cells through their target migration domain. Similar to the control, the total glia extension of CD8-GFP tagged membranes spread evenly to occupy the posterior region of the photoreceptor differentiation domain (Fig. 1D-F). In contrast, this adaptability was not observed in the non-glial cells of the eye disc, as *ey*-Gal4 driven expression of Rho1 RNAi caused severe reductions in disc size, apoptosis/cell death and interference with photoreceptor differentiation (Fig. S1). Therefore, we investigated the mechanism for the compensatory response in glia. Glia nuclear area, defined by the expression of the glia-specific transcription factor Repo, was significantly increased upon Rho1 knockdown (Fig. 1D’-F’). Furthermore, glial cell ploidy was also highly increased to an average of 15n, ranging from 8n to 25n (Fig. 1H). In control discs only the two subperineurial glial cells were significantly polyploid (10n), as previously described (Silies et al., 2007; Unhavaithaya and Orr-Weaver, 2012; Von Stetina et al., 2018; Zulbahar et al., 2018). This ability of Rho1 depleted glial cells to compensate for a reduced number and sustain a normal domain of association to neurons is not a general phenomenon. A similar phenotype has been observed upon overexpression of the E3 ubiquitin ligase Fizzy related (Fzr) protein in perineurial glia (Silies et al., 2007). Rho1 knockdown resulted in a significant decrease in viability, mainly after pupation, during pupal stages (Fig. 1I), revealing a crucial role for Rho1 expression in glia.

**Fig. 1.**
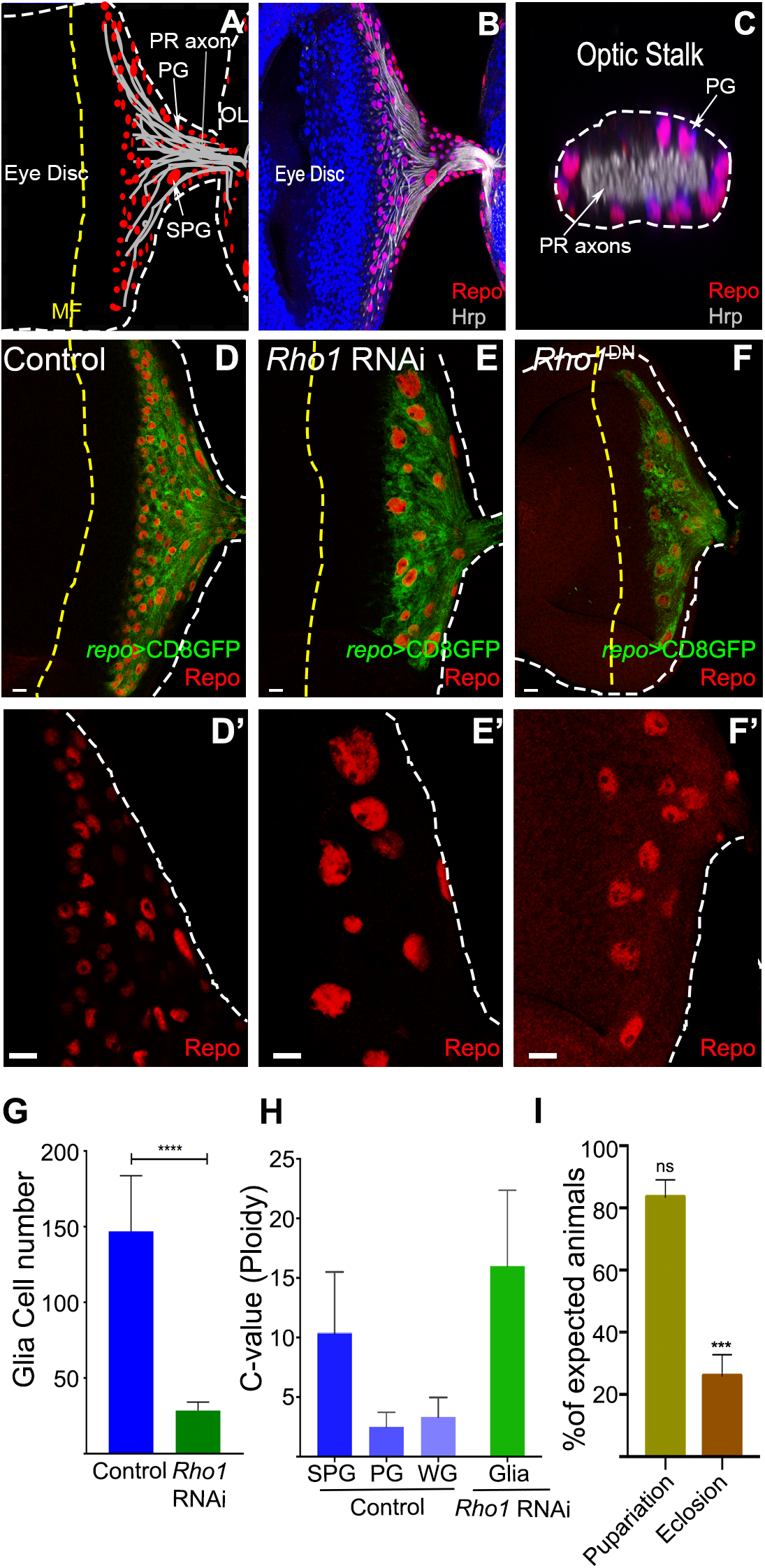
Rho1 *loss-of-func*tion in glia causes a switch from mitosis to endoreplication. A) Schematic representation of the *Drosophila* eye imaginal disc (ED) and optic lobe (OL) indicating the different glial cells observed, namely perineurial glia (PG) and subperineurial glia (SPG). White dashed line surrounds the eye disc and OL. Yellow dashed line represents the morphogenetic furrow (MF). Gray lines represent photoreceptor (PR) axons. (B) Control ED showing glia staining by Repo (red) and PR axons by Hrp (grey). (C) Control transversal cut of the OS showing PG nuclei surrounding PR axons. (D) Control (*repo*4.3>CD8GFP>*LacZ*), (E) *Rho1* RNAi (*repo*4.3>CD8GFP>*Rho1* RNAi) and (F) *Rho1* dominant-negative (DN; repo4.3>CD8GFP>Rho1^DN^). Glia membranes (Repo>CD8GFP) are shown in green. D’ – F’ are magnification of glia staining with Repo (red). Scale bars correspond to 10 μm. (G) Graph showing glial cell number in Control (*repo*>*Dcr-2*>*LacZ*, n=19) and *Rho1* RNAi (*repo*>*Dcr-2*>*Rho1* RNAi, n=10) from eye discs with 7 to 12 Photoreceptor rows (PhR). p<0.0001. (H) Graph representing retinal glia C-value (ploidy) in Control (*repo*>*Dcr-2*>LacZ, n=6) and *Rho1* RNAi (*repo*>*Dcr-2*>*Rho1* RNAi, n=10) from eye discs with 7 to 12 PhR. (I) Developmental viability of *repo*>*Rho1* RNAi animals. Percentage of expected pupariation and eclosion. Results are mean ± s.e.m. (***p < 0.01; n.s., not significant).

We next aimed to identify the pathway of Rho1 upstream regulators and downstream effectors required to prevent polyploidization of eye disc glial cells. Rho guanine nucleotide exchange factors (RhoGEFs) regulate the activation of Rho family members. Pebble (Pbl), the fly ortholog of vertebrate ECT2, is essential for the formation of a contractile ring and the initiation of cytokinesis, acting at the cell cortex as a Rho-specific RhoGEF (Prokopenko et al., 1999). Human Ect2 also activates RhoA at the onset of mitosis to induce the actomyosin remodelling that drives both mitotic rounding and cortical stiffening (Matthews et al., 2012; Rosa et al., 2015). Interestingly, in a similar manner to Rho1 knockdown, expression of a UAS-*pebble(pbl)* RNAi construct with *repo*-Gal4 led to a reduction in number and polyploidization of glial cells in the eye disc (Fig. 2A, compare with Control Fig. 1D). In contrast, knocking down RhoGEF2, a regulator of cell shape changes during *Drosophila* embryogenesis (Azevedo et al., 2011; Hacker and Perrimon, 1998), or the single *Drosophila* Rho GDP dissociation inhibitor (RhoGDI)(Golding et al., 2019) did not reveal proliferation/polyploidization defects in glial cells (Fig. 2B, C). Phenotypes for all the genes tested in this study were confirmed by using at least a second nonoverlapping RNAi line. Pebble (RhoGEF)/Rho GTPase control several pathways during cytokinesis including the recruitment of the formin Diaphanous to the equatorial cortex to stimulate F-actin assembly (Castrillon and Wasserman, 1994), and the activation of Rok/ROCK (Rho-associated protein kinase) and Citron Kinase, which activate non-muscle myosin II (Ishizaki et al., 1996; Mizuno et al., 1999; Yamashiro et al., 2003). Interestingly, we observe that anillin a scaffolding protein, encoded by *scra*, which links contractile elements within the furrow to the plasma membrane and to microtubules (Echard et al., 2004; Hickson and O’Farrell, 2008; Piekny and Glotzer, 2008) was required for glial ploidy control (Fig. 2D), while ROCK was not (Fig. 2E, F). The latter result is in accord with a more crucial role for Citron Kinase than for ROCK during cytokinesis (Madaule et al., 1998). Overall, we showed that the Pebble (RhoGEF)/Rho1/Anillin pathway regulates the ploidy level of retinal glia in the eye disc.

**Fig. 2.**
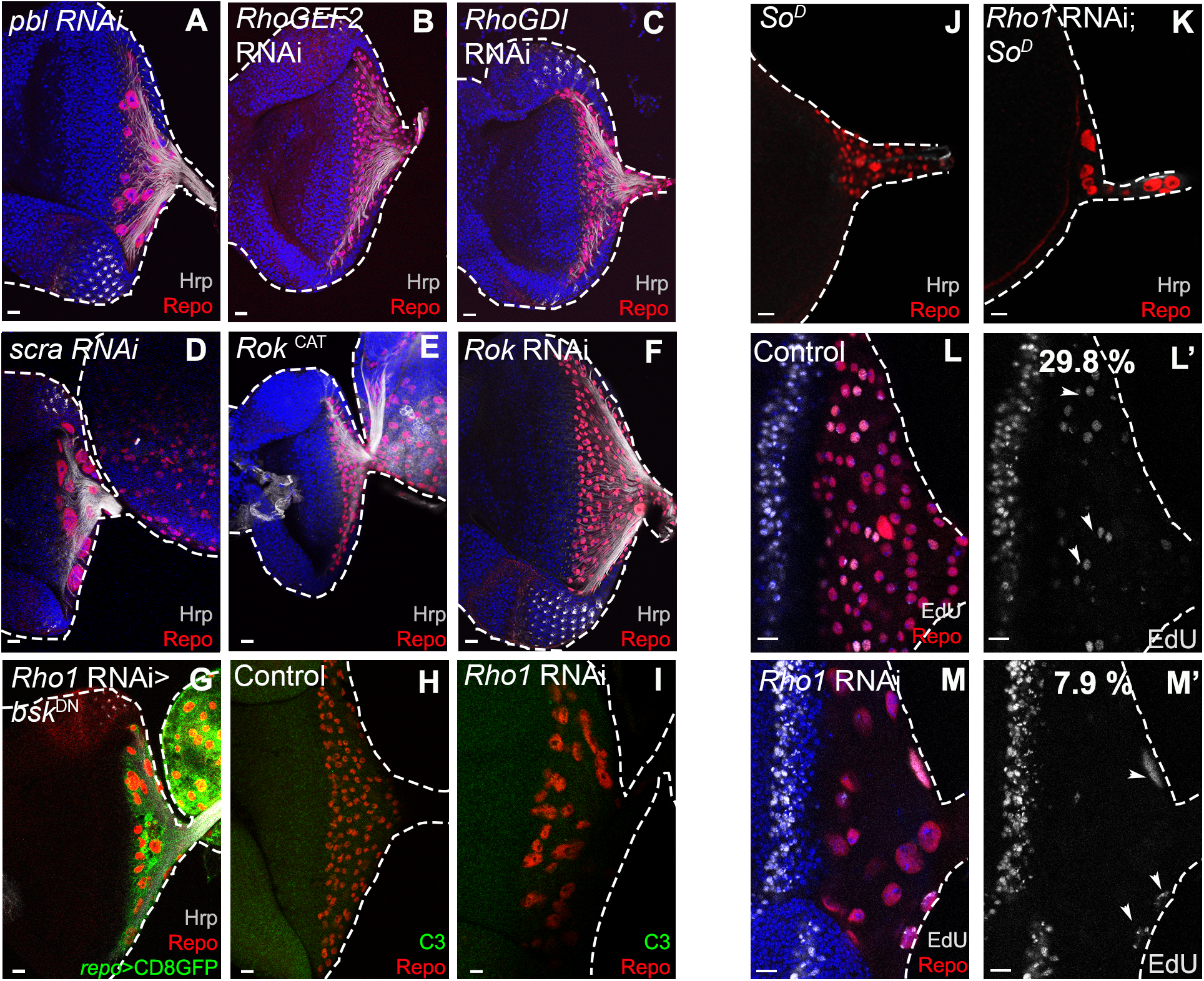
The Pebble/Rho1/Anillin pathway regulates the glia ploidy level in the eye disc. (A) *pbl* RNAi (*repo*4.3>CD8GFP>*pbl* RNAi) (B) *RhoGEF2* RNAi (*repo*4.3>CD8GFP>*RhoGEF2* RNAi) (C) *RhoGDI* RNAi (*repo*4.3>CD8GFP>*RhoGDI* RNAi) (D) *scra* RNAi (*anillin*; *repo*>*Dcr-2*>*scra* RNAi) (E) Constitutively-active Rok, *Rok*^CAT^ (*repo*4.3>CD8GFP>*Rok*^CAT7.1^) (F) *Rok* RNAi (*repo*4.3>CD8GFP>*Rok* RNAi). (G) Co-expression of *bsk*^DN^ does not alter *repo*>*Rho1* RNAi phenotype (*repo*4.3>CD8GFP>*Rho1* RNAi>*bsk*^DN^). Green corresponds to glial cell membranes (CD8GFP). (H - I) Cleaved Caspase-3 (C3) staining (green) in (H) Control (*repo*>*Dcr-2*>LacZ) and (I) *Rho1* RNAi (*repo*>*Dcr-2*>*Rho1* RNAi). (J - K) Rho1 represses polyploidization in the absence of photoreceptor differentiation. Dominant-negative *sine oculis* (*so*^D^) (J) and *repo*4.3>*Rho1* RNAi1 in *so*^D^ background (K). Hrp in shown in grey. EdU staining (grey) in (L) Control (*repo*>*Dcr-2*>LacZ) and (M) *Rho1* RNAi (*repo*>*Dcr-2*>*Rho1* RNAi). (L’-M’) show EdU staining in grey. Arrowheads point towards EdU staining in retinal glia. % of EdU positive retinal glia are indicated (n=6 in control and n=8 in *Rho1* RNAi ED with 7-11 PhR; **p=0.0027). Repo staining (glia) is shown in red. Hrp is shown in grey, except for L - M (corresponds to EdU). White dashed line surrounds the eye disc. DAPI stains DNA in blue. Scale bars correspond to 10 μm.

### Regulation of the glia-specific Rho1 knockdown phenotype during eye development

While the formation of polyploid cells has been implicated in genomic instability and cancer (Davoli and de Lange, 2011), it is also a physiological mechanism associated to the differentiation of some cell types, such as megakaryocytes that undergo endomitosis as a failure of late cytokinesis caused by defects in the contractile ring and Rho GTPase signalling (Gao et al., 2012; Lordier et al., 2008). In the highly proliferative *Drosophila* wing disc, cytokinesis failure induced by knockdown of the Septin peanut (pnut) caused the formation of tetraploid cells and massive apoptosis that was dependent on Jun N-terminal kinase (JNK) activity (Gerlach et al., 2018). Interestingly, in Rho1-knockdown glial cells the resulting increase in ploidy is not significantly rescued by the co-expression of a dominant-negative version of JNK (bsk^DN^; Fig. 2G), and it does not lead to apoptosis, evaluated by Caspase3 activation (Fig. 2H-I). Furthermore, blocking any possible early apoptosis by expressing the apoptotic inhibitor p35 failed to prevent glial polyploidization in Rho1-knockdown eye discs (Fig. S2 A, B). This further supports that some cell types such as retinal glial cells tolerate Rho1 knockdown or cytokinesis failure better than wing or eye imaginal disc cells (Fig. S1). Interestingly, in a dominant-negative *sine oculis* (*so*^D^) genetic background, where photoreceptor differentiation is, as expected, absent or reduced to one photoreceptor row in 90% of eye discs (Fig. 2 J, K) (Kenyon et al., 2005; Roederer et al., 2005), glial cells depleted for Rho1 failed to enter the disc proper, but these stalled cells in the optic stalk already displayed some increase in ploidy (Fig. 2 J, K). We next evaluated the contribution of proliferation defects to the reduced number of glial cells in *repo*>*Rho1* RNAi eye discs. In comparison to controls, we detected reduced incorporation of 5-Ethynyl-2’-deoxyuridine (EdU) by Rho1-knockdown glial cells in the eye disc (Fig. 2 L, M). On one hand this suggests that the increase in ploidy also occurs in retinal glial cells after their migration from the optic stalk. This is also supported by the detection of pH3-positive large polyploid cells that progressed to mitosis (Fig. S2 C, D), revealing that at least a fraction of the cells undergoes an endomitotic type of endoreplication cell cycle. On the other hand, the resulting polyploid cells have diminished DNA replication (Fig. 2L, M), so the function of other factors could be at play to modulate cell proliferation and genome instability. Results presented here suggest that the initial stage of polyploidization is directly caused by Rho1 loss-of-function, independently of glia interactions with photoreceptors, and it allows for further glial polyploidization in the disc to accommodate for a proper matching of photoreceptors and glia surface area.

### Rho1 functions in perineurial glia to inhibit eye disc glia polyploidization

To further understand which glia subtypes require Rho1 function to control eye disc glia number and ploidy, we analysed Rho1 expression in the different glia layers. We used an anti-Rho1 antibody (Magie et al., 2002)(Fig. S3A, B), and a GFP trap inserted in the Rho1 gene (encoding *Rho1*-*GFP*; Fig. S3C). The expression patterns obtained with both approaches were very similar. Rho1 was detected in subperineurial glial cells and in perineurial glia. Next, we induced Rho1 inhibition using glia-subtype specific drivers. When dominant-negative Rho1^DN^ is expressed under the control of a subperineurial-specific driver, (*moody*-GAL4) (Sasse et al., 2015; Schwabe et al., 2005), no significant change in the ploidy of glial cells in the eye disc was observed (Fig. 3A, B). However, when Rho1^DN^ was expressed specifically in perineurial glia using the C527 driver (Hummel et al., 2002), the main features of pan-glial Rho1 depletion were recapitulated. Mainly, the number of glia in the eye disc was markedly reduced and its nuclear size significantly increased, both in the disc and in the optic stalk region (Fig. 3C, D, arrowheads). Interestingly, the glia nuclear size in the optic lobes was less affected (Fig. 3C, D, asterisks). Wrapping glia is the third glia subtype identified in the eye imaginal disc. When Rho1 was depleted using a wrapping glia-specific driver (Mz97-GAL4)(Hummel et al., 2002), we detected no significant differences in cell numbers of total glia and differentiated wrapping glia, nor in nuclear sizes (Fig. 3E-G). Thus, Rho1 is not required for the maintenance of a differentiated and diploid status in wrapping glia. To exclude the possibility that using the Mz97-GAL4 driver significant inhibition of Rho1 is only achieved after cells are already committed to differentiation, we evaluated differentiation upon earlier and pan-glia Rho1 repression using *repo*-Gal4 and the *sprouty*-*lacZ* differentiation marker (Sieglitz et al., 2013). Interestingly, we detected expression of *sprouty*-*lacZ* in a large subset of the polyploid glial cells induced by Rho1 knockdown (Fig. 3H, I). In fact, the percentage of differentiated wrapping glial (over the total repo positive cells) increases from 49 % in control to 69 % in Rho1 RNAi (Fig. 3J). Thus, our results suggest that Rho1 knockdown in perineurial glia induces polyploidization, but subsequently Rho1 is not required for the commitment of polyploid cells to the wrapping glia fate.

**Fig. 3.**
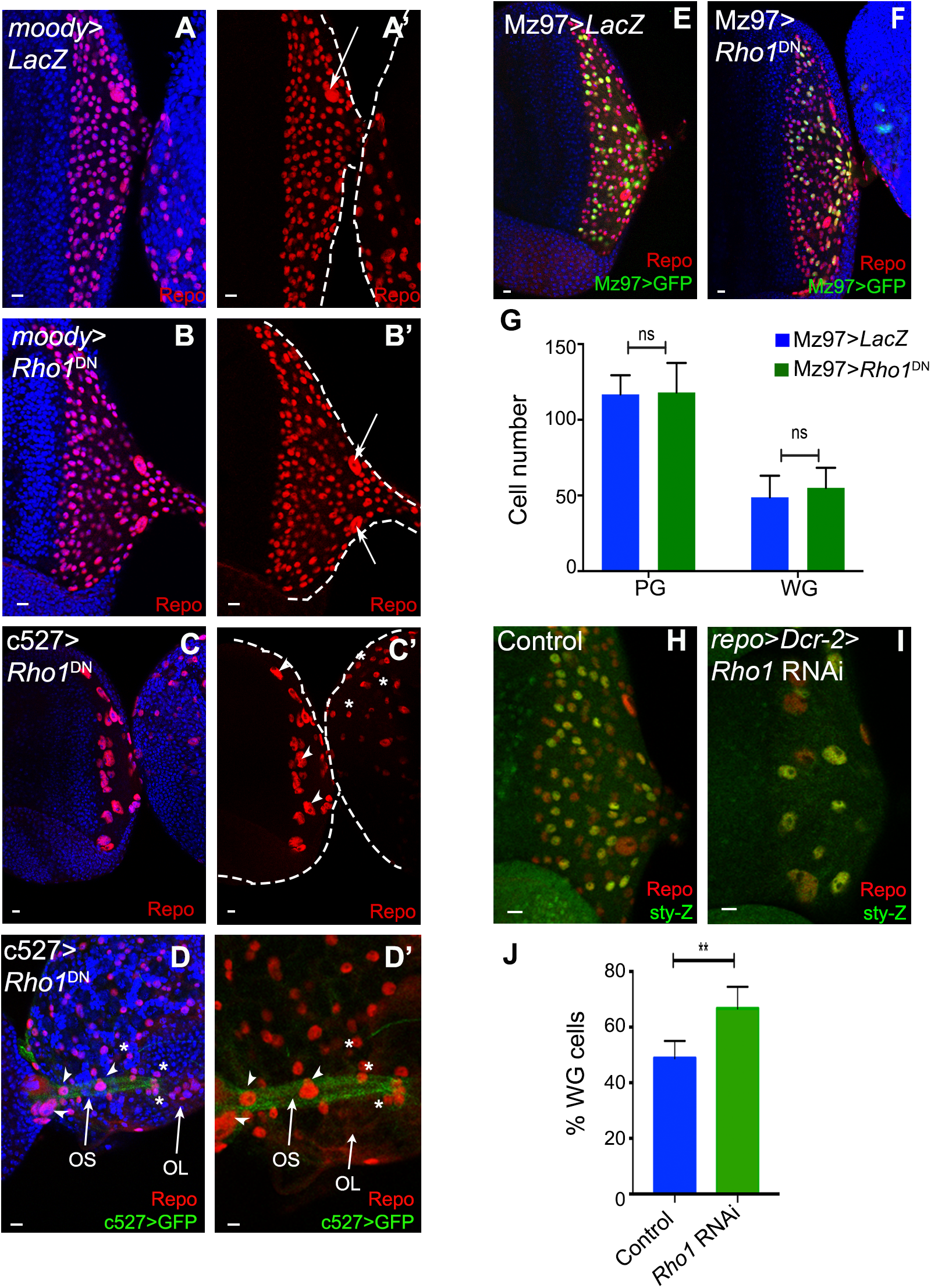
Rho1 functions in perineurial glia to inhibit eye-disc glia polyploidization. A - B) *Rho1* LOF in SPG (B, *moody*>*Rho1*^DN^) show a similar phenotype as control (A, *moody*>LacZ). Arrows points to SPG nuclei. (C) PG *Rho1*^DN^ (c527>*Rho1*^DN^) demonstrates a phenotype similar with pan glia *Rho1* LOF. Nuclear size is increased in the ED (arrowheads) while the glia nuclear size (asterisk) in OL remains similar to control (A’). A’ – C’ shows glia in red. (D) c527>P2A-GFP>*Rho1*^DN^ showing differences in glia nuclear size between the eye optic stalk (OS; arrows) and the OL (asterisk). D’ shows glia in red and PG membranes in green. (E, F) Wrapping glia (WG; Mz97 expressing cells) are shown in green in Control (E; Mz97>Stinger>LacZ) and *Rho1*^DN^ (F; Mz97>Stinger>*Rho1*^DN^). (G) Graph representing PG and WG cell numbers in Control (Mz97>Stinger>LacZ) and *Rho1*^DN^ (Mz97>Stinger>*Rho1*^DN^). Average of 5 eye imaginal discs. (H - I) WG are labelled by β-galactosidase from *sprouty*-LacZ (*sty*-Z; green) in (H) control (*repo*>*Dcr-2*>*sty*-Z) and (I) *Rho1* RNAi (*repo*>*Dcr-2*>*sty*-Z>*Rho1* RNAi #29002). (J) Graph showing the percentage of WG (sty-positive) in Control and Rho1 RNAi (genotypes as H and I). Average of 10 (control) and 6 ED (*Rho1* RNAi). ** p=0.0016. Glia is stained by Repo in red. DNA was counterstained with DAPI (blue). Scale bars correspond to 10 μm.

### Rho1-depleted glia initiate axonal wrapping but fail to complete ensheathment of 8-axons fascicles

In control eye discs, differentiated wrapping glia contact and initiate wrapping of nascent photoreceptor axons (Silies et al., 2007; Tavares et al., 2015). However, when we expressed Rho1 RNAi we observed a decrease in the total number of wrapping glia nuclei in the eye disc and in the optic stalk (Fig. 4A, B). To evaluate if axonal wrapping is regulated by Rho1, we first analyzed glia membranes through the expression of the membrane marker CD8-GFP. This analysis showed that in the eye disc Rho1-depleted glia is still capable of extending membrane projections to initiate axonal wrapping in a similar fashion to control, with axons projecting to the optic stalk (Fig. 4C, D). Further analysis of the optic stalk showed that photoreceptor axons are less physically insulated from hemolymph in Rho1 RNAi, being closer to the Laminin-rich neural lamella, due to the reduction in glial nuclei and membrane extension (Fig. 4E-J). By transmission electron microscopy (TEM) it is possible to visualize two glial cell layers between the photoreceptor axons and the ECM corresponding to the subperineurial cells and the more external perineurial glia (Fig. 4K). In Rho1 RNAi optic stalks only one glial cell layer was visible in some regions (Fig. 4L). Interestingly, while in the controls most clusters of eight ommatidial axons are wrapped as one fascicle (Silies et al., 2007; Tavares et al., 2015), in Rho1 RNAi we observe fewer wrapping glia processes and a significant reduction in the ensheathment of 8-axons ommatidial fascicles (Fig. 4M, N; Fig. S4). The decreased fasciculation is also observed by immunofluorescence, where axonal staining appears less condensed (Fig. 4I, J).

**Fig. 4.**
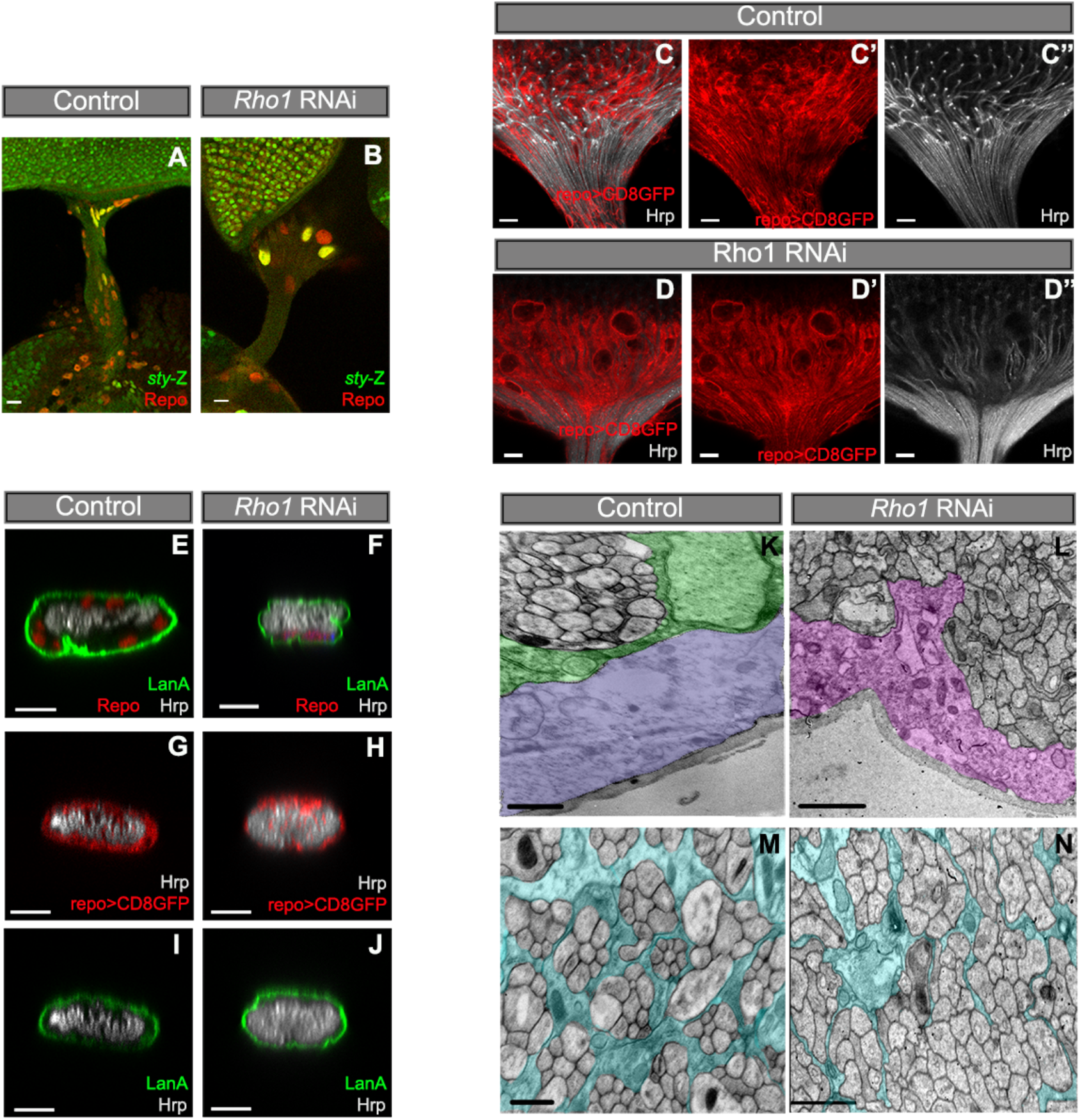
Rho1 knockdown causes wrapping defects in the optic stalk. A - B) OS of (A) control (*repo*>*Dcr-2*>*sty*-Z) and (B) *Rho1* RNAi (*repo*>*Dcr-2*>*sty*-Z>*Rho1* RNAi #29002) labelled with β-galactosidase to detect *sprouty*-LacZ (*sty*-Z; green). C) Control (*repo*4.3>CD8GFP>LacZ) and (D) *Rho1* RNAi (*repo*4.3>CD8GFP>*Rho1* RNAi). Repo>CD8GFP is shown in red (C’ and D’) and PR axons are stained by Hrp and shown in grey (C’’ and D’’). (E, G and I) Control (Repo>Dcr-2>LacZ) and (F, H and J) *Rho1* RNAi (*repo*>*Dcr-2*>*Rho1* RNAi) transversal cut of the OS. Laminin A (LanA) is shown in green and glia nuclei (Repo) and membranes (Repo>CD8GFP) in red. Hrp stains PR axons in grey. Transversal cut of the OS analysis by TEM from (K and M) Control (*repo*>*Dcr-2*>LacZ) and (L and N) *Rho1* RNAi (*repo*>*Dcr-2*>*Rho1* RNAi). Wrapping glia (blue) enwraps each R1-R8 ommatidia laying apical to SPG (green). PG cells (lilac) form a layer externally to the SPG. In *Rho1* RNAi instead of PG and SPG layer, only a single glia layer is visible (SPG-like in pink). Glia (repo) is shown in red. (A – J) Scale bars correspond to 10 μM. (K – N) Scale bars correspond to 1 μm.

Overall, our results show that Pebble (RhoGEF)-Rho1-Anillin are specifically required in perineurial glia to limit polyploidization, and that glial cells display high plasticity and adaptability to the polyploid status being able to differentiate and to initiate photoreceptor axonal wrapping. However, Rho1 function is necessary for the later ensheathment of 8-axons fascicles and viability, which is a question that will be interesting to approach in future work.

## Material and Methods

### Fly husbandry

Most crosses were raised at 25 °C under standard conditions. The following stocks (described in FlyBase, unless stated otherwise) were used: *repo*-Gal4 , c527-Gal4 (Hummel et al., 2002); *moody*-Gal4 (Sasse et al., 2015; Schwabe et al., 2005), *repo*4.3>CD8GFP (*moody*-Gal4 and *repo*4.3-Gal4 were gifts from Christian Klämbt, University of Münster); UAS-*Dcr-2*, Mz97-GAL4,UAS-Stinger (BDSC_#9488) (Hummel et al., 2002), Rho1-GFP (ZCL1957) UAS-*Rho1*^DN19,1.3^ (BDSC_#7327) (Barrett et al., 1997), UAS-*Rho1* RNAi (BDSC_#29002 in Fig. 3 H-I, elsewhere: BDSC_#27727), UAS-*pebble* (#109305 and #35349), UAS-*RhoGEF2* RNAi (#34643, #31239, #10577), UAS-*RhoGDI* RNAi (46155, #105765), UAS-*scra* RNAi (#53358), UAS-Rok RNAi (#3793, #104675, #3797), UAS-*Rok*^CAT^ (#6668, #6669), UAS-lacZ, UAS-*yki*^ACT^ (#28817), UAS-*dMyc*, UAS-*bsk*^DN^K53R (#9311), *sprouty*-LacZ [54], w1118, UAS-CD8GFP, UAS-CD4tdTOM, *so*^D^/CyO (#4287). RNAis were validated with all the lines described above.

For viability experiments, crosses between *repo*-Gal4/TM6B and UAS-*Rho1* RNAi or a neutral UAS-lacZ were setup, and vials were scored daily for pupariation and eclosion according to genotype. Pupariation and eclosion were expressed as percentage of expected values, after normalisation: the proportion of animals (*repo*-Gal4 > UAS*-Rho1* RNAi)/(UAS-Rhoi/TM6B) was divided by the (*repo*-Gal4 > UAS*-lacZ*)/(UAS-lacZ/TM6B) control ratio and converted in percentage of expected pupariation and eclosion. Statistical comparison of pupariation and eclosion rates between Rho1 and lacZ conditions was done using unpaired t-test with Welch correction. The experiments were repeated at least thrice.

### Immunohistochemistry

Eye-antennal imaginal discs were dissected in cold Phosphate Buffer Saline (PBS) and fixed in 3.7% formaldehyde/PBS for 20 minutes. Immunostaining was performed using standard protocols (Marinho et al., 2013). Primary antibodies used were: mouse anti-repo antibody at 1:5 (8D12 anti-Repo, Developmental Studies Hybridoma Bank, DSHB), rat anti-repo at 1:1000 (a gift from Dr. Benjamin Altenhein, Institut für Genetik, Germany), rabbit anti-repo at 1:2000 (a gift from Dr. Benjamin Altenhein), rabbit anti-pH3 antibody at 1:1000 (Upstate), rat anti-Elav 1:100 (7E8A10 DSHB), goat anti-HRP antibody Cy5-conjugated at 1:100 (Jackson ImmunoResearch), rabbit anti-cleaved Caspase-3 at 1:200 (9661, Cell Signaling), rabbit anti-β-galactosidase antibody 1:2000 (Cappel, 55976, MP Biomedicals), rabbit anti-Laminin A antibody at 1:200 (a gift from Herwig O. Gutzeit) and mouse anti-Rho1 at 1:100 (p1D9, DSHB). Appropriate Alexa Fluor conjugated secondary antibodies used were from Molecular Probes. For Ethynyl deoxyuridine (EdU) experiments, dissected eye-antennal imaginal discs were incubated in 10 μM EdU/PBS for 10 minutes, at room temperature, washed with PBS and fixed as described above (Eusebio et al., 2018). Alexa Fluor Azide detection was performed according to Click-iT EdU Fluor Imaging Kit (Invitrogen).

Image stack of eye discs were obtained with the Leica SP5 confocal system and processed with Adobe Photoshop. Glia quantification was performed on the ventral most single plane encompassing the highest number of glia nuclei. Results presented here are representative examples from samples with more than 6 eye imaginal discs. Glial cells were counted, glia-covered eye disc area and glia nuclear area measured in Fiji. Mean and standard deviation were calculated for each case using GraphPad Prism.

### Ploidy quantification

DAPI quantification protocol was described before (Zulbahar et al., 2018). DAPI staining was at 100 ng/ ml DAPI in 1× PBS containing 0.1% Triton X-100 for 2 h at room temperature. DAPI fluorescence intensity was measured using Fiji software. The corrected total integrated density (CTCF) was calculated for each nucleus using the following function: CTCF=integrated density – (area of selected nucleus × mean fluorescence of background readings). Background readings were made by measuring the fluorescence three times in regions in which no nuclei were present. Ploidy was then calculated by normalizing each glia (repo positive) nucleus to the average of 10 Elav-positive diploid photoreceptors, imaged on the same eye disc with the same laser settings.

### Statistical analysis

To determine the statistical significance of the glial cell number, glia nuclear area, and C-value comparisons in Fig. 1 we applied a Mann–Whitney test. Statistical significance of the comparisons of perineurial glia and wrapping glia numbers between Control and Rho1 RNAi in Fig. 3G was done using 2-way ANOVA. Comparison of wrapping glia cell number between Control and Rho1 RNAi in Fig. 3J was done using Mann Whitney test. GraphPad Prism was used, p-value *=p<0.05; **=p<0.01; ***=p<0.001; ****=p<0.0001. Error bars present in all graphs represent the standard deviation.

### Electronic Microscopy

Electronic microscopy imaging was performed as previously reported (Marinho et al., 2011; Martins et al., 2017; Tavares et al., 2015). For evaluation of ultrastructural features, electron micrographs were examined from 3 different animals per genotype.

## Acknowledgements

We thank Christian Klämbt, Herwig O. Gutzeit, Benjamin Altenhein, the Bloomington Drosophila Stock Center, the Vienna Drosophila RNAi Center, the Drosophila Genetic Resource Center, and the Developmental Studies Hybridoma Bank for reagents. We thank Emiliana Pereira and Pedro Silva for technical assistance. The authors acknowledge the support of the Advanced Light Microscopy (ALM) and Histology and Electron Microscopy (HEMS) i3S Scientific Platforms, members of the national infrastructure PPBI - Portuguese Platform of Bioimaging (PPBI-POCI-01-0145-FEDER-022122). This work was funded by Norte-01-0145-FEDER-000008 - Porto Neurosciences and Neurologic Disease Research Initiative at I3S and Norte-01-0145-FEDER-000029 - Advancing Cancer Research: From basic knowledge to application, both supported by Norte Portugal Regional Operational Programme (NORTE 2020), under the PORTUGAL 2020 Partnership Agreement, through the European Regional Development Fund (FEDER). LT was supported by an FCT Postdoc Fellowship (SFRH/BPD/95336/2013).

