## Supplementary Materials for "The Pebble/Rho1/Anillin pathway controls polyploidization and axonal wrapping activity in the glial cells of the *Drosophila* eye"

**Supp. Figure Legends**

**Fig. S1. Rho1 *loss-of-func*tion (LOF) in the eye imaginal disc**

Ey>Drc-2>Rho1 RNAi ED show almost absence of differentiation, in comparison with Control. Glia is stained by Repo (red) and photoreceptor (PR) axons by Hrp (grey). Inlet (dashed line) shows pyknotic (apoptotic) nuclei (arrows). DAPI stains DNA in blue. Scale bars correspond to 10 μm.

**Fig. S2. *Rho1*-depleted polyploid retinal glia initiate mitosis independently of apoptosis**

(A, B) No difference observed in eye discs in (A) Control (*repo*>*Dcr-2*>LacZ) and (B) *Rho1* RNAi (*repo*>*Dcr-2*>*Rho1* RNAi>p35) for Cleaved Caspase-3 (C3) staining (green) and Repo (red). (C, D) Presence of Phospho-Histone H3 (pH3) staining (green) in (C) Control (repo>Dcr-2>LacZ) and (D) Rho1 RNAi (repo>Dcr-2>Rho1 RNAi). Glia is stained by Repo (red) and photoreceptor (PR) axons by Hrp (grey). DAPI stains DNA in blue. Scale bars correspond to 10 μm.

**Fig. S3. *Rho1* expression in glial cells**

A) Rho1 expression detected by an α-Rho1 antibody (green) in retinal glia. PG membranes can be identified by c527>CD4tdTOM in grey. Glia nuclei are shown in red. Rho1 staining is visible in the outermost glia membrane corresponding to perineurial glia (arrows). B) Rho1 expression detected by an α-Rho1 antibody (green) in SPG. SPG membranes can be identified by moody>CD8GFP in red. PG expression is also visible (arrow). C) Rho1-GFP trap line showing Rho1 expression (green) in the outermost glia layer of the optic stalk corresponding to PG. Glia is stained by Repo (red). Scale bars correspond to 10 μm.

**Fig. S4. Wrapping deficiencies in Rho1 RNAi optic stalk.**

Transversal cut of the OS analysis by TEM from Control (*repo*>*Dcr-2*>LacZ) and *Rho1* RNAi (*repo*>*Dcr-2*>*Rho1* RNAi). Wrapping glia (blue) enwraps each R1-R8 ommatidia in Control but large regions of unwrapped axons are found in Rho1 RNAi.

### Fig. S1

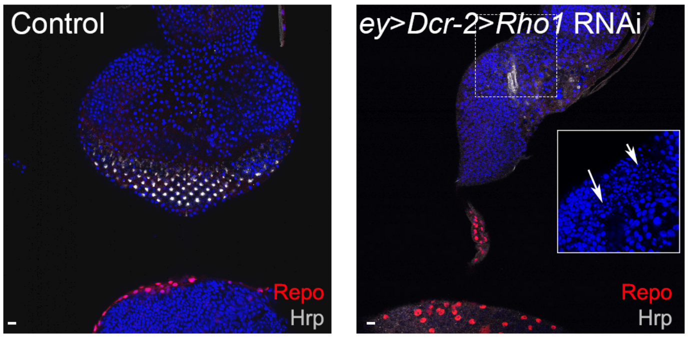

**Fig. S2**

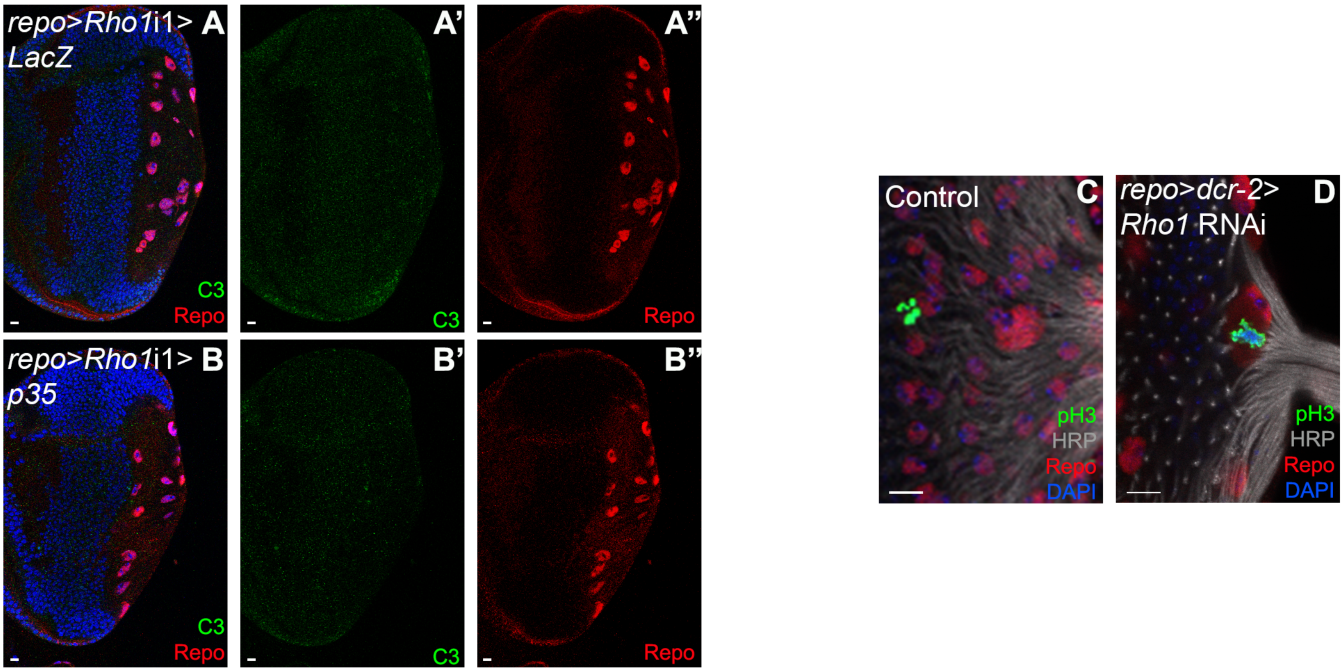
repo>Rho1i1> LacZ
A
A'
A''
C3
Repo
C3
Repo
repo>Rho1i1> p35
B
B'
B''
C3
Repo
C3
Repo
Control
C
pH3
HRP
Repo
DAPI
repo>dcr-2> Rho1 RNAi
D
pH3
HRP
Repo
DAPI

### Fig. S3

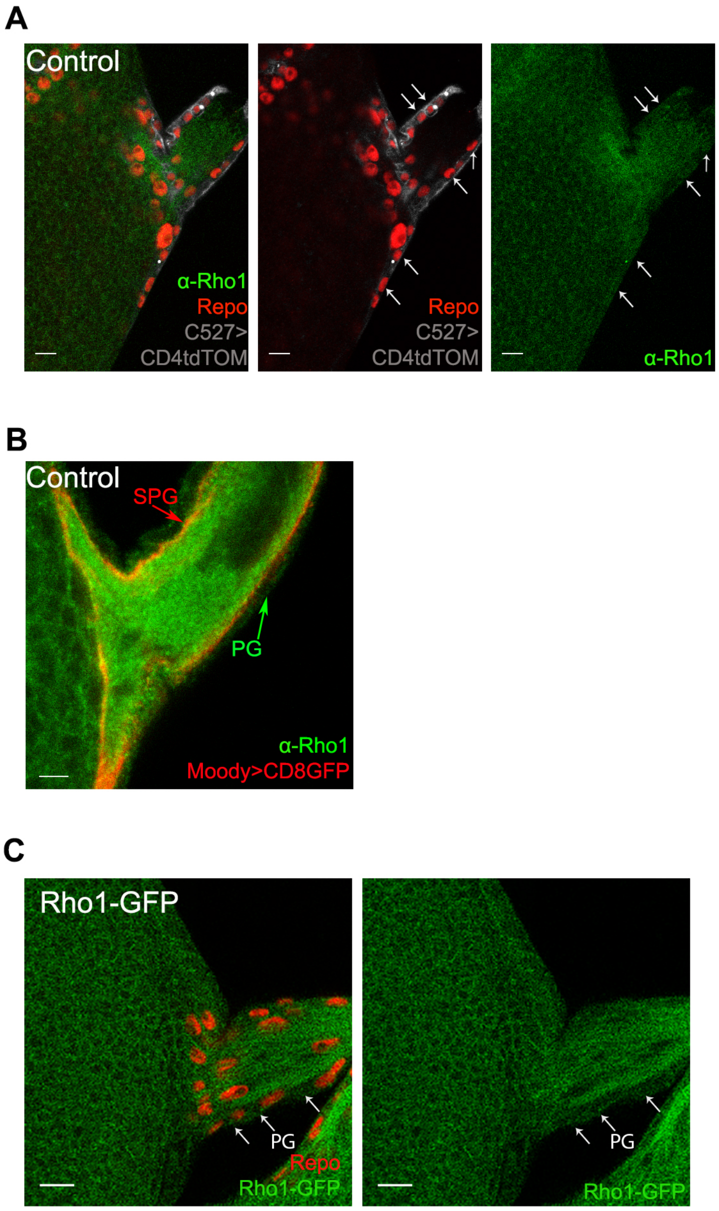
A
Control
α-Rho1
Repo
C527>
CD4tdTOM
Repo
C527>
CD4tdTOM
α-Rho1
B
Control
SPG
PG
α-Rho1
Moody>CD8GFP
C
Rho1-GFP
PG
Repo
Rho1-GFP
PG
Rho1-GFP

**Fig. S4**

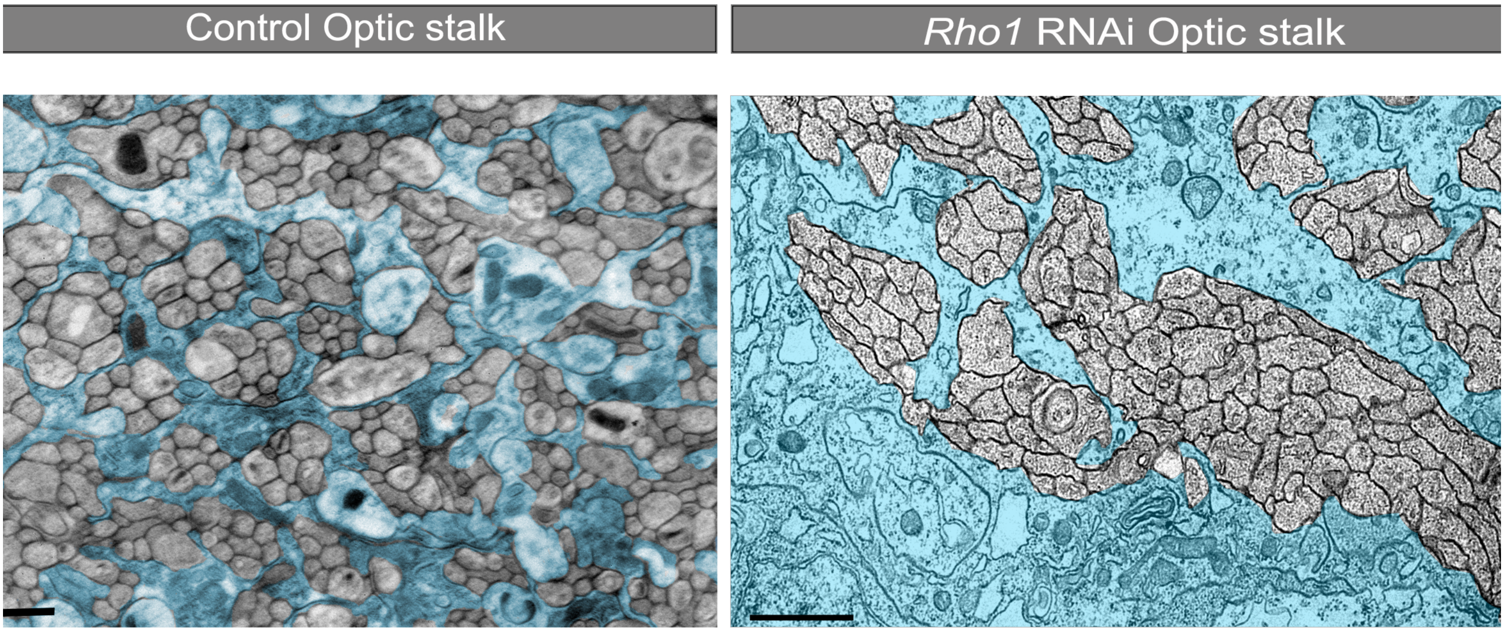
